# Identification of a simple and novel cut-point based CSF and MRI signature for predicting Alzheimer’s disease progression that reinforces the 2018 NIA-AA research framework

**DOI:** 10.1101/443325

**Authors:** Priya Devanarayan, Viswanath Devanarayan, Daniel A. Llano, the Alzheimer’s Disease Neuroimaging Initiative (ADNI)

## Abstract

The 2018 NIA-AA research framework proposes a classification system with beta-<u>A</u>myloid deposition, pathologic <u>T</u>au, and <u>n</u>eurodegeneration (ATN) for the diagnosis and staging of Alzheimer’s Disease (AD). Data from the ADNI (AD neuroimaging initiative) database can be utilized to identify diagnostic signatures for predicting AD progression, and to determine the utility of this NIA-AA research framework. Profiles of 320 peptides from baseline cerebrospinal fluid (CSF) samples of 287 normal, mild cognitive impairment (MCI) and AD subjects followed over a 3-10 year period were measured via multiple reaction monitoring (MRM) mass spectrometry. CSF Aβ_42_, total-Tau (tTau), phosphorylated-Tau (pTau-181) and hippocampal volume were also measured. From these candidate markers, optimal diagnostic signatures with decision thresholds to separate AD and normal subjects were first identified via unbiased regression and tree-based algorithms. The best performing signature determined via cross-validation was then tested in an independent group of MCI subjects to predict future progression. This multivariate analysis yielded a simple diagnostic signature comprising CSF pTau-181 to Aβ_42_ ratio, MRI hippocampal volume and a novel PTPRN peptide, with a decision threshold on each marker. When applied to a separate MCI group at baseline, subjects meeting this signature criteria experience 4.3-fold faster progression to AD compared to a 2.2-fold faster progression using only conventional markers. This novel 4-marker signature represents an advance over the current diagnostics based on widely used marker, and is much easier to use in practice than recently published complex signatures. In addition, this signature reinforces the ATN construct from the 2018 NIA-AA research framework.

**Disclosures:** Viswanath Devanarayan is an employee of Charles River Laboratories, and as such owns equity in, receives salary and other compensation from Charles River Laboratories.

Data collection and sharing for this project was funded by the Alzheimer’s Disease Neuroimaging Initiative (ADNI) (National Institutes of Health Grant U01 AG024904) and DOD ADNI (Department of Defense award number W81XWH-12-2-0012). ADNI is funded by the National Institute on Aging, the National Institute of Biomedical Imaging and Bioengineering, and through generous contributions from the following: AbbVie, Alzheimer’s Association; Alzheimer’s Drug Discovery Foundation; Araclon Biotech; BioClinica, Inc.;Biogen; Bristol-Myers Squibb Company; CereSpir, Inc.; Eisai Inc.; Elan Pharmaceuticals, Inc.; Eli Lilly and Company; EuroImmun; F. Hoffmann-La Roche Ltd and its affiliated company Genentech, Inc.; Fujirebio; GE Healthcare; IXICO Ltd.; Janssen Alzheimer Immunotherapy Research & Development, LLC.; Johnson & Johnson Pharmaceutical Research & Development LLC.; Lumosity; Lundbeck; Merck & Co., Inc.; Meso Scale Diagnostics, LLC.; NeuroRx Research; Neurotrack Technologies; Novartis Pharmaceuticals Corporation; Pfizer Inc.; Piramal Imaging; Servier; Takeda Pharmaceutical Company; and Transition Therapeutics. The Canadian Institutes of Health Research is providing funds to support ADNI clinical sites in Canada. Private sector contributions are facilitated by the Foundation for the National Institutes of Health (www.fnih.org). The grantee organization is the Northern California Institute for Research and Education, and the study is coordinated by the Alzheimer’s Disease Cooperative Study at the University of California, San Diego. ADNI data are disseminated by the Laboratory for Neuro Imaging at the University of Southern California.

## Introduction

Tools to provide an early diagnosis and prediction of progression to Alzheimer Disease (AD) are of critical importance. Early diagnosis allows caregivers to plan for additional needs which will decrease the overall financial burden of the illness [1, 2]. In addition, early diagnosis may help identify common comorbidities such as depression or undernutrition [3, 4], and may spur lifestyle interventions to mitigate some of the cognitive impairments associated with aging [5]. Finally, identifying individuals who are more likely to progress will help enrich clinical trial populations with subjects with more rapid progression, potentially shortening trial duration.

Early pathological changes of AD are seen years before the clinical diagnosis of AD. Most studies have shown that individuals with mild cognitive impairment (MCI) carry AD pathological burden and have a substantial risk (~10-15% per year) of development of dementia [6, 7]. Thus, as new therapeutics are developed that target AD-related pathology, MCI may represent a state during which early intervention may change the trajectory of patient outcomes. However, therapeutics targeting Aβ will likely carry potential risks of significant side-effects, as documented in clinical trials [8-10], thus limiting their use to those with a high risk of subsequent cognitive decline. Therefore, what is needed is an approach to accurately identify MCI patients with the highest risk of conversion to AD.

Multiple potential biomarkers have been identified to aid in the prediction of conversion of MCI to AD. For example, cognitive and behavioral biomarkers have been proposed to identify individuals at high-risk for conversion [11-13]. In addition, biomarkers based on brain imaging or measurements in bodily fluids have been identified (for recent reviews see [14, 15]). The latter groups of biomarkers have recently been organized into a generalizable research framework. This framework, labeled AT(N), describes three classes of biomarkers: 1) “A” or aggregated amyloid-based (e.g., cerebrospinal fluid (CSF) Aβ_42_ levels, amyloid positron emission tomography (PET)), 2) “T” or aggregated tau-based (e.g., CSF phosphorylated tau [pTau-181], tau PET) and 3) “N” or neuronal degeneration-based (e.g., volumetric magnetic resonance imaging (MRI), fluorodeoxyglucose (FDG) PET, CSF total tau (tTau)) [16, 17]. Furthermore, Jack et al, 2018, advocate extending this to the ATX(N) framework, where X can include additional markers from the multiarray–omics platforms. This research framework is intended to form a common approach by which investigators can communicate about and classify novel biomarkers, thereby allowing their more rapid integration into current research.

CSF-based biomarkers have been of interest since they represent an assessment of biochemical changes in the central nervous system. The most commonly-observed changes in the CSF of AD subjects have been a reduction of Aβ_42_ and increase in pTau-181 [18, 19]. We recently identified a 16-analyte CSF signature which showed higher sensitivity and specificity than any combination of Aβ_42_, tTau and pTau-181 for the diagnosis of AD vs. controls, and, when applied to an independent dataset of MCI subjects, outperformed traditional biomarkers in prediction of conversion to AD [20]. Unfortunately, a complicated 16-analyte signature is not practical for clinical purposes. Multi-analyte signatures require quality-control measures for each analyte and do not provide an intuitive understanding of how changes in the biomarker impact the disease. Therefore, here, using data from only the AD and age-matched Normal (NL) subjects from the ADNI (AD neuroimaging initiative) database, we identify optimal diagnostic signatures with decision thresholds on a few markers that separate AD and NL subjects via unbiased regression and tree-based algorithms [21, 22]. The best performing signatures from the NL and AD subjects were then tested in an independent group of MCI subjects at baseline to determine their ability to predict the future progression. By developing a simple, yet powerful, signature for prediction of MCI to AD, this work confirms and extends the AT(N) framework and provides a potential new tool for clinicians to use to advise decision making and for researchers to enrich clinical trials with MCI subjects with a higher likelihood of conversion.

## Methods

Data used for this research were mostly identical to that used in Llano et al (2017), except that the progression data on MCI now extends to another two years, and we include data from the conventional biomarkers (CSF amyloid/tau and MRI HV) in the analysis. For the sake of completeness, we repeat some of the key information pertaining to these data in this paper. The ADNI database (adni.loni.usc.edu) utilized in this research was launched in 2003 as a public-private partnership, led by Principal Investigator Michael W. Weiner, MD. The primary goal of ADNI has been to test whether serial MRI, PET, other biological markers, and clinical and neuropsychological assessments can be combined to measure the progression of MCI and early AD. For up-to-date information, see www.adni-info.org. This study was conducted across multiple clinical sites and was approved by the Institutional Review Boards of all of the participating institutions. Informed written consent was obtained from all participants at each site. Data used for the analyses presented here were accessed on February 24, 2018. Although the ADNI database continues to be updated on an ongoing basis, most newly added biomarker data are from later time points (i.e., beyond 1 year), in contrast to the baseline data used in this study.

### Patient Population

This research was focused on only those subjects in the ADNI database for whom data from both the conventional markers (CSF amyloid/tau and MRI HV) and novel markers (320 peptides from the multiple reaction monitoring (MRM) proteomics panel) were available at baseline. This included 287 subjects with AD, MCI and NL from the ADNI study that received clinical, neuropsychological, and biomarker assessments which were repeated every six months for a period of 3 to 10 years. NL individuals were free of memory complaints or depression and had a Mini-Mental State Examination (MMSE) score above 25 and a Clinical Dementia Rating (CDR) score of 0. MCI individuals could have MMSE scores of 23 to 30 and required a CDR of 0.5 and an informant-verified memory complaint substantiated by abnormal education-adjusted scores on the Wechsler Memory Scale Revised—Logical Memory II. AD patients could have MMSE scores of 20 to 27 and a CDR of 0.5 or 1.0.

### Imaging

All participants received 1.5 Tesla (T) structural MRI at baseline and at every six months for the next several years. In addition, approximately 25% also received 3.0 T MRI. Cognitive assessments and neuroimaging procedures were carried out within two weeks of each other. In this research, we utilized only the baseline HV data measured via MRI and computed using the FreeSurfer software at the University of California in San Francisco. Details regarding this software can be found in the “UCSF FreeSurfer Methods” PDF document under “MR Image Analysis” in the ADNI section of https://ida.loni.usc.edu/) as well as in [23-25].

### CSF Samples

CSF levels of Aβ_42_, tTau, and pTau-181 were determined using Innogenetics’ INNO-BIA AlzBio3 immunoassay on a Luminex xMAP platform (see [19] for details). These CSF samples were also processed in the Caprion Proteomics platform that uses mass spectrometry to evaluate the ability of a panel of peptides to discriminate between disease states and predict disease progression. The CSF multiplex MRM panel was developed by Caprion Proteomics in collaboration with the ADNI Biomarker Consortium Project Team. A total of 320 peptides generated from tryptic digests of 143 proteins were used in this study (see Supplemental Table 1 and supplemental table in [20] for list of peptides and proteins).

Details regarding the technology, quality control and validation of the MRM platform can be found in the Use of Targeted Mass Spectrometry Proteomic Strategies to Identify CSF-Based Biomarkers in Alzheimer’s Disease Data Primer (found under Biomarkers Consortium CSF Proteomics MRM Data Primer at ida.loni.usc.edu). In brief, as described in the data primer and in [26], plasma proteins were depleted from CSF samples using a Multiple Affinity Removal System (MARS-14) column, and digested with trypsin (1:25 protease:protein ratio). The samples were then lyophilized, desalted and analyzed by LC/MRM-MS analysis on a QTRAP 5500 LC-MS/MS system at Caprion Proteomics. MRM experiments were performed on triple quadrupole (Q) mass spectrometers. The first (Q1) and third (Q3) mass analyzer were used to isolate a peptide ion and a corresponding fragment ion. The fragment ions were generated in Q2 by collision induced dissociation. All peptide levels are presented as normalized and log2-transformed intensities as we and others have done previously [20, 26], which is identical to the manner in which they were provided in the quality-controlled dataset.

### Statistical Methods

Optimal combinatorial signatures with simple decision thresholds on each marker were first identified from the AD and NL subjects. This was performed in an unbiased, data driven manner via regression and tree-based computational algorithms called Patient Rule Induction Method [21] and Sequential BATTing [22]. Prior to application of these algorithms on the 320-peptide MRM panel, promising peptide candidates were identified using the same algorithm utilized in [20], using the logistic regression model with Lasso penalty [27] and a bootstrap procedure [28] to improve the stability of the lasso parameter estimate.

The predictive performance of the optimal signature from each algorithm for differentiating the AD and NL subjects was then evaluated via 10 iterations of five-fold cross-validation. In this procedure, the original data were divided into five random subsets (folds), each fold was left out one at a time, and the remaining four folds were used to derive a signature, which was then used to predict the disease state of each subject in the left-out fold. This process was carried out for each left-out fold one at a time and the predictions of all the five left-out folds were aggregated. For better stability and robustness, this cross-validation procedure was repeated 10 times and the median of each these performance measures was calculated. All steps of the model building and signature derivation process were fully embedded within this cross-validation to further reduce any possible bias [29].

The optimal signature from the best performing algorithm determined via the above cross-validation procedure (i.e., the signature that best differentiated AD and NL subjects) was then tested on a separate independent group of 135 MCI subjects at baseline, to predict their future progression to AD. MCI subjects predicted to be AD-like (“Signature Positive” at baseline) were considered as future converters to AD, and those predicted to be NL-like (“Signature Negative” at baseline) were considered as non-converters. These baseline predictions were then compared to the follow-up clinical data. Performance metrics such as the PPV, NPV and overall accuracy were calculated by comparing the predictions to the known progression status of the MCI subjects to AD over the next 36 months. Comparisons of the performance metrics between different signatures were carried out via exact McNemar’s test.

The performance of this signature was then evaluated in terms of its ability to differentiate the future time to progression from MCI to AD of these baseline signature-positive and signature-negative MCI subjects via Kaplan-Meier analysis. For this evaluation, the progression of MCI subjects to AD over the entire future time course until the last follow-up visit was taken into consideration.

This analysis procedure was carried out separately for the following subsets of markers, along with APOE genetic status, age, gender and education (4 markers):

- MRI brain HV: 5 total markers (the 4 markers above + HV)
- CSF Aβ_42_, tTau, pTau-181, ratios of tTau to Aβ_42_ & pTau-181 to Aβ_42_ (AT): 9 total markers
- AT + HV: 10 total markers, and
- AT + HV + 320 peptides from the CSF MRM panel: 330 total markers

This evaluation of the AD versus NL peptide signature on the future progression of a separate group of MCI subjects to AD not only served as an independent verification of the utility of the signature, but also put it to a greater test to see whether it is robust enough to address a different and more important question related to the prediction of future progression of the MCI subjects to AD. The analysis procedure described here is summarized in Figure 1.

**Figure 1:**
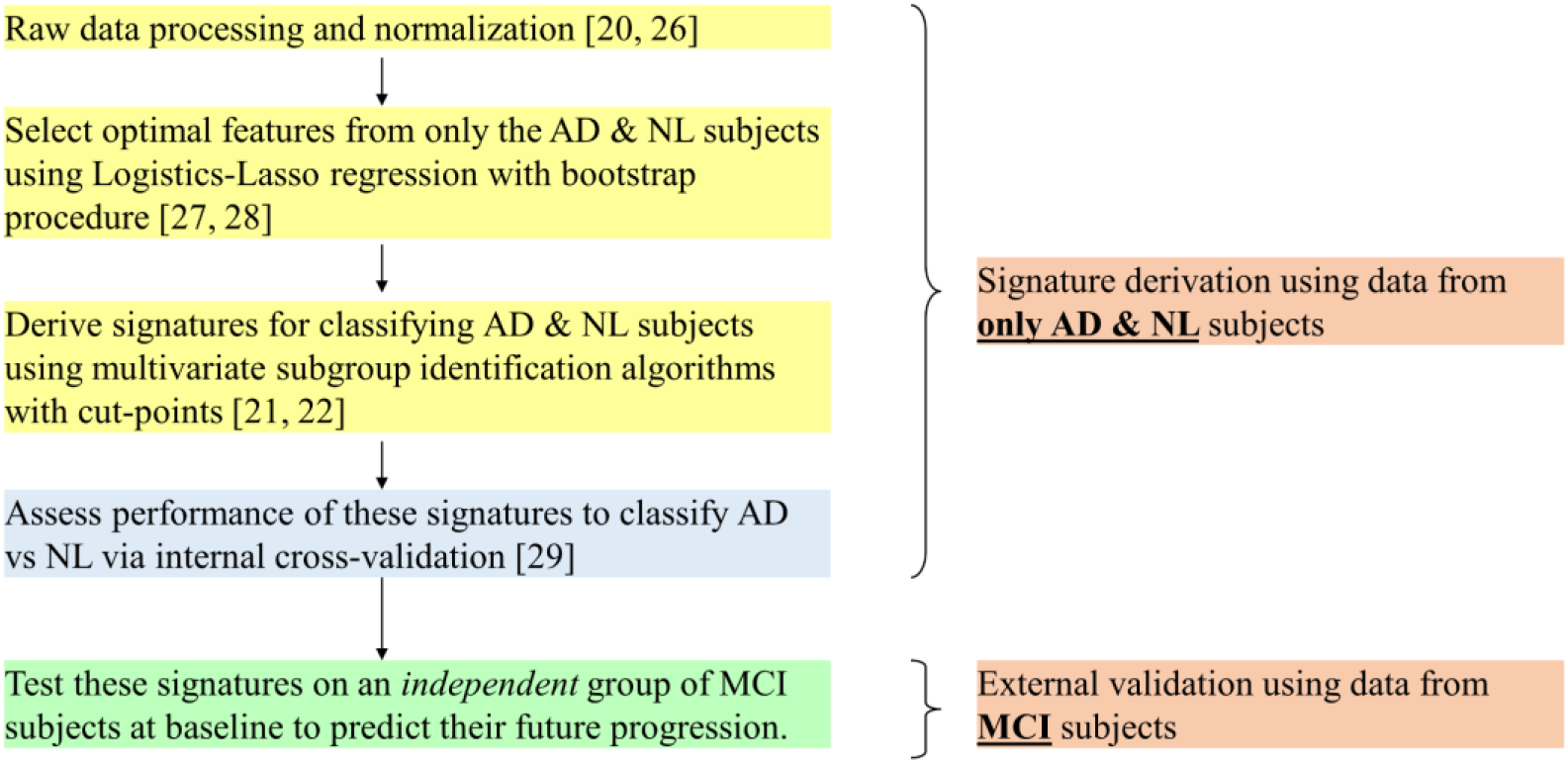
Statistical analysis flow-scheme

All analyses related to predictive modeling and signature derivation were carried out using R (http://www.R-project.org), version 3.4.1, with the publicly available package, SubgrpID [22]. The time to progression analysis of the derived signatures and related assessments were carried out using JMP®, version 13.2.

## Results

### Disease-state Demographics

Table 1A summarizes the key demographics of the 66 AD, 135 MCI and 86 NL subjects, and Table 1B provides a breakdown of the 135 MCI subjects in terms of their future progression. The subjects were balanced across groups in terms of age and education (both p>0.05). There were significantly more males (59.1%) than females (40.9%) in the study, though similar numbers of male and female MCI subjects converted to AD over a three-year period (44% vs. 54.6%, p=0.248, Chi-squared test). As shown previously [30], the presence of at least one copy of the APO-E4 allele was a risk factor for AD (71.2% AD, 52.6% MCI and 24.4% NL, p < 0.0001, Chi-squared test). In addition, this allele also tracked with MCI to AD progression over a 36-month period (37.5% of non-E4 vs. 56.3% of E4 progressed to AD, p=0.029, Chi-squared test).

**Table 1:** Disease-state demographics

|  |  | **AD** | **MCI** | **NL** |
| --- | --- | --- | --- | --- |
|  |  | **(n=66)** | **(n=135)** | **(n=86)** |
| **Gender (n)** | **M** | 37 | 91 | 44 |
|  | **F** | 29 | 44 | 42 |
| **Apo-E (n)** | **E4** | 47 | 71 | 21 |
|  | **Non-E4** | 19 | 6 | 65 |
| **Age**<br>(years, Mean +/- SD) |  | 75.1 +/- 7.5 | 74.8 +/- 7.4 | 75.8 +/- 5.6 |
| **Education**<br>(years, Mean +/- SD) |  | 15.1 +/- 3 | 16 +/- 3 | 15.6 +/- 3 |
| **MMSE**<br>(Mean +/- SD) |  | 23.5 +/- 1.9 | 26.9 +/- 1.7 | 29.1 +/- 1 |

|  |  | **MCI to AD converters** | **Stable MCI** |
| --- | --- | --- | --- |
|  |  | **(n=64)** | **(n=71)** |
| **Gender (n)** | **M** | 40 | 51 |
|  | **F** | 24 | 20 |
| **Apo-E (n)** | **E4** | 40 | 31 |
|  | **Non-E4** | 24 | 40 |
| **Age**<br>(years, Mean +/- SD) |  | 74.9 +/- 7.6 | 74.7 +/- 7.2 |
| **Education**<br>(years, Mean +/- SD) |  | 15.6 +/- 3.0 | 16.4 +/- 2.9 |
| **MMSE**<br>(Mean +/- SD) |  | 26.4 +/- 1.7 | 27.4 +/- 1.6 |

### Disease state classification

The distribution of the conventional biomarkers for NL, MCI and AD subjects are shown in Figures 2A-D. While the means significantly differ across groups (p<0.0001), the considerable overlap of expression levels greatly limits the diagnostic utility of any of these markers on their own.

**Figure 2:**
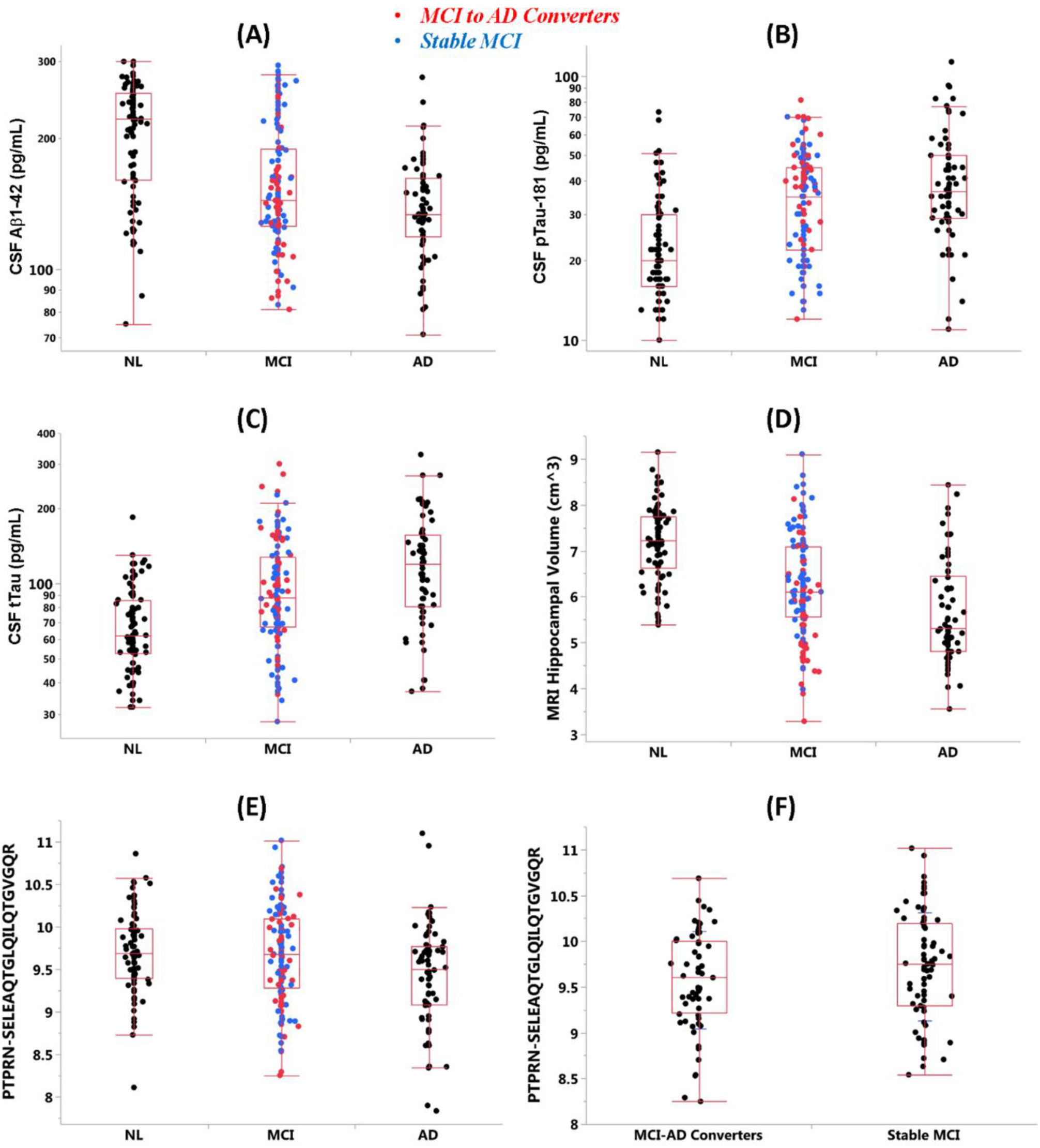
Distribution of four conventional markers of AD (A: CSF Aβ_42_, B: CSF pTau-181, C: CSF tTau, D: MRI HV) are shown for the NL, MCI and AD subjects at baseline. Among the MCI subjects, those that progressed to AD over 36 months are shown in red and rest are shown in blue. The bottom and top ends of the box represent the first and third quartiles respectively, with the line inside the box representing the median. Lines extending out of the ends of the box indicate the range of the data, minus the outliers. The points outside the lines are the low and high outliers. E) Distribution of PTPRN.SELE peptide (in normalized log2 transformed intensity units) is shown across the NL, MCI and AD groups at baseline, and F) for the baseline MCI subjects that either progressed to AD or remained stable over the next 36 months.

Multivariate analysis of the various markers using data-driven computational algorithms described above yielded optimal signatures for differentiating the disease states and prediction of disease progression. These signatures are summarized in Table 2. Interestingly, the signature derived from the conventional and novel markers took a very simple form based on only a few markers, with representations from both the conventional markers and the novel MRM panel, along with a cut-point on each of them; it took the form of HV < 7.65 cm^3^, ratio of pTau to Aβ_42_ > 0.09 and a PTPRN peptide (sequence SELEAQTGLQILQTGVGQR, referred to here as PTPRN.SELE) < 10.22 intensity units. Figures 2E-F show the significant decline of PTPRN.SELE in AD relative to both NL (p=0.002) and MCI (p=0.004), and a trend towards a decline in baseline MCI subjects that progress to AD in the future (p=0.065).

**Table 2:** Performance of optimal signatures

| Data type | Diagnostic Criteria for Signature positive | AD versus Normal Diagnosis (internal cross-validation) |  |  | 36m MCI Progression to AD (independent validation) |  |  |
| --- | --- | --- | --- | --- | --- | --- | --- |
|  |  | PPV | NPV | Accuracy | PPV (MCI to AD) | NPV (Stable MCI) | Accuracy |
| **AT** | $t\text{Tau} / \text{Ab1-42} > 0.59$ | 71.6% | 80.5% | 76.5% | 58.1% | 66.1% | 61.7% |
| **HV** | $\text{HV} < 6.41$ and ApoE4 + | 92.7% | 74.8% | 79.6% | 61.2% | 60.5% | 60.7% |
| **AT + HV** | $\text{HV} < 7.0$, $p\text{Tau} > 18.1$, and $t\text{Tau} / \text{Ab1-42} > 0.36$ | 73.4% | 78.4% | 76.3% | 60.2% | 70.2% | 64.4% |
| **AT + HV + MRM** | $\text{HV} < 7.65$, $p\text{Tau} / \text{Ab1-42} > 0.09$, and $\text{PTPRN.SELE} < 10.22$ | **75%** | **79.6%** | **77.6%** | **61.6%** | **77.6%** | **67.4%** |

### Prediction of MCI to AD progression

For disease state classification, the signatures derived from all data scenarios have similar levels of overall accuracy, with no discernable advantage of adding novel markers from the MRM panel to the conventional markers. However, for the prediction of 36-month progression in the independent group of 135 MCI subjects at baseline, the signature derived from the collection of both conventional and novel markers significantly outperforms the signatures based on the conventional markers (p=0.0002), with the NPV increasing from 70.2% to 77.6% (p=0.0032) and the PPV increasing slightly from 60.2% to 61.6% (p=0.0107). Thus, the addition of a novel PTPRN peptide from the MRM panel to the conventional AD markers substantially improves the prediction of 36-month disease progression in MCI subjects at baseline.

Based on the available 3-10 year follow-up clinical data available on these subjects, the performance of the optimal signatures from all the scenarios was further assessed on this independent group of baseline MCI subjects with respect to their future time to progression. Table 3 includes a summary of the 25^th^ percentile, median, and 75^th^ percentile time to progression of the signature negative and signature positive subjects, and the overall Hazard Ratio with 95% confidence bands. Based on these results, the optimal combination of conventional markers showed a Hazard Ratio of 2.2 suggesting that the MCI subjects meeting the criteria of this signature experience 2.2-fold faster progression to AD. However, the MCI subjects that meet the signature criterion from the scenario that includes the PTPRN peptide experience 4.3-fold faster progression to AD, as shown in Figure 3. To determine if the impact of PTPRN was likely isolated to the particular peptide fragment (PPRN.SELE, the other two PTPRN peptides (AEAPALFSR, referred to as PTPRN.AEAP and LAAVLAGYGVELR, referred to as PTPRN.LAAV) in the MRM panel were also assessed. The pairwise correlations between these three peptides are all over 87% (data not shown).

**Figure 3:**
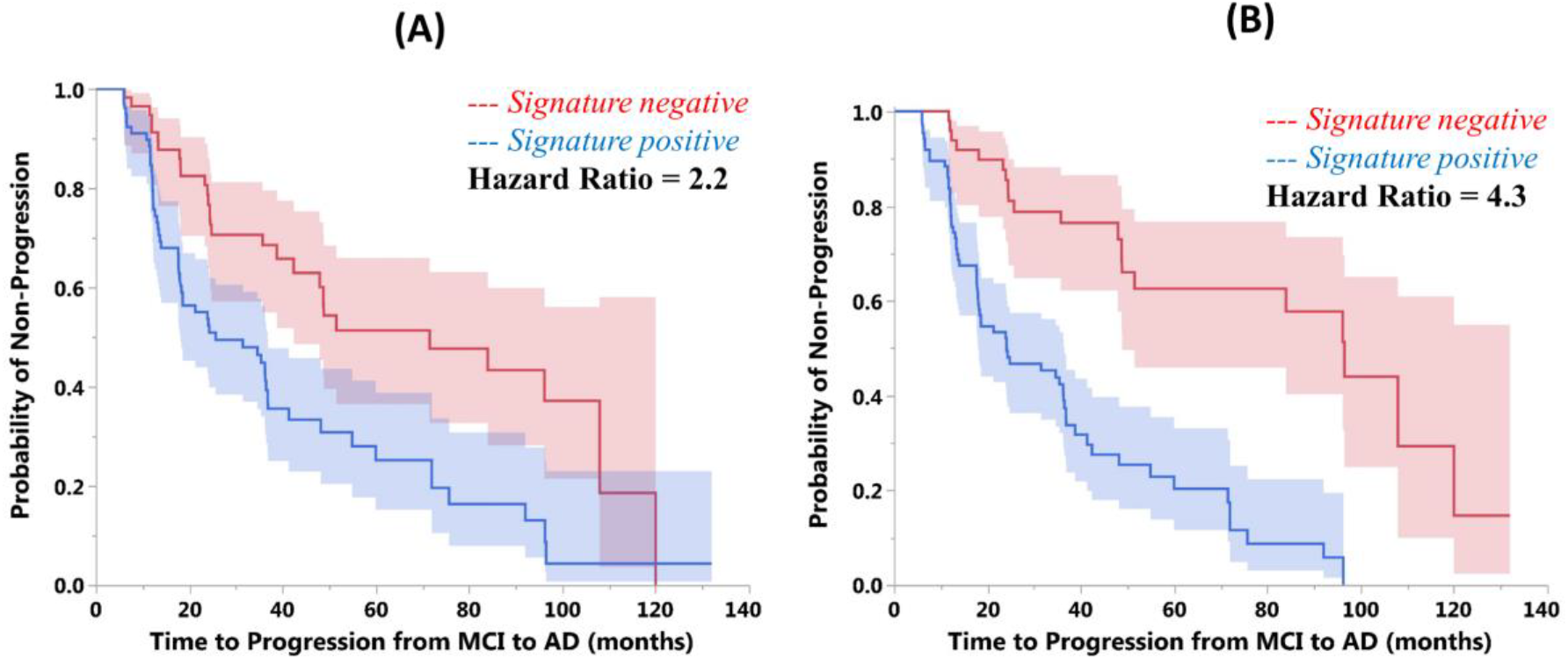
Time to progression profiles of the signature positive versus signature negative MCI subjects with the shaded 95% confidence bands are shown here via Kaplan-Meier analysis. The effect of signature based on only the conventional markers (HV and AT) is illustrated in Figure 4A and the signature with both the conventional markers and the novel PTPRN.SELE peptide from the MRM panel is shown in Figure 3B. Patients meeting the signature criterion that includes this PTPRN peptide experience 4.3-fold faster progression to AD (hazard ratio = 4.4), relative to the 2.2-fold faster progression without this peptide.

**Table 3:** Time to progression (T2P) of MCI subjects to AD using optimal signatures

| Data type | Diagnostic Criteria for Signature positive | Signature Negative |  | Signature Positive |  | Hazard Ratio (95% C.I.) |
| --- | --- | --- | --- | --- | --- | --- |
|  |  | N | T2P (months)<br>Q1, Q2, Q3 | N | T2P (months)<br>Q1, Q2, Q3 |  |
| **AT** | $t\text{Tau} / \text{Ab1-42} > 0.59$ | 59 | 23.4, 71.6, 108 | 76 | 13.6, 25.7, 72.0 | 1.9 (1.2, 3.1) |
| **HV** | $\text{HV} < 6.41$ and ApoE4 + | 86 | 18.6, 48.2, 108 | 49 | 13.1, 31.5, 60.0 | 2.0 (1.3, 3.2) |
| **AT + HV** | $\text{HV} < 7.0$, $p\text{Tau} > 18.1$,<br>and<br>$t\text{Tau} / \text{Ab1-42} > 0.36$ | 57 | 24.4, 71.6, 108 | 78 | 12.6, 25.7, 72.0 | 2.2 (1.4, 3.6) |
| **AT + HV + MRM** | $\text{HV} < 7.65$,<br>$p\text{Tau} / \text{Ab1-42} > 0.09$, and<br>$\text{PTPRN.SELE} < 10.22$ | 49 | 48.0, 96.5, 120 | 86 | 12.7, 24.1, 54.9 | 4.3 (2.5, 7.7) |

## Discussion

### Summary

We examined the ability of a simple optimized multivariate signature comprising conventional biomarkers combined with an array of novel CSF peptides from the ADNI database to both classify AD disease state and to predict MCI to AD conversion. We observed that both conventional AD biomarkers (HV and CSF pTau/Aβ_42_ ratio) and conventional biomarkers combined with an array of novel CSF peptides performed similarly in terms of classifying disease state (AD vs. NL). However, when these optimized signatures were applied to an independent group of MCI subjects, the signature combining conventional markers with a novel peptide analyte derived from PTPRN substantially outperformed the conventional biomarkers in predicting MCI to AD conversion by nearly twofold. In addition, the combined signature contains only four elements: HV, CSF Aβ_42_, tTau and the PTPRN.SELE peptide, thus making it simple enough to be tractable for clinical and research purposes. These data may also open new lines of investigation regarding the role of PTPRN in AD as well as confirming and extending the proposed AT(N) framework for AD biomarkers.

#### PTPRN and AD

PTPRN is expressed widely in neurons throughout the mouse and human brain, including areas associated with AD neurodegeneration such as hippocampus and neocortex [31, 32]. It is a membrane-spanning protein phosphatase with cytoplasmic and luminal components and is found in the membranes of secretory granules. The gene for PTPRN is also highly expressed in pancreatic islet cells, and antibodies against this protein are found in type 1 diabetes, hence its alternative name islet-antigen 2 [33]. Deficiency in PTPRN is associated with glucose intolerance in animal models [34] as well as impaired learning [35]. Given the associations between diabetes, insulin resistance and AD [36-38], it is possible that the PTPRN/AD association seen in the current study points to a new and specific role for metabolic dysregulation in the pathophysiology of AD to complement other metabolic hypotheses of AD [39, 40].

Several previous studies have identified PTPRN as a potential marker of AD. For example, downregulation of the expression of the PTPRN gene has been observed in the hippocampus of sporadic AD subjects [41] as well as the posterior cingulate area of early-onset AD and presenilin-1 mutation-related dementia [42]. In addition, when incorporated into a three-gene classifier, PTPRN expression levels have been found to discriminate between patients with AD pathology and no symptoms, and those with only AD pathology [43]. Finally, in a preliminary study of genetic interactions with CSF pTau levels for predicting MCI to AD conversion, PTPRN levels showed differences with respect to CSF pTau levels in MCI to AD converters compared to non-converters [44].

#### Implications of the prediction of MCI-AD conversion

Over the years, several groups have examined the ability of multi-modal combination biomarkers (i.e., combinations of imaging, cognitive, body fluid and other markers) to predict the conversion of MCI to AD. Ideally, utilizing an approach such as the AT(N) framework, a combination biomarker should merge several orthogonal measurements reflecting different underlying biological processes. Larger combinations of biomarkers have the potential to increase the predictive power of the combination biomarker. The multiplicity of biomarkers is limited by clinical reality such that it is often impractical and costly to obtain multiple studies in individual patients. Therefore, a challenge in developing combination biomarkers is to develop combinations that provide high predictive MCI to AD accuracy and are clinically feasible.

Here, we have identified a 4-marker signature that combines volumetric MRI and CSF testing, both feasible clinical tests, that outperforms standard biomarkers in the prediction of MCI to AD. Although other studies have found that combinations of volumetric MRI and CSF measures can predict MCI to AD conversion [45-50], a unique aspect of the current biomarker signature is that it was initially developed using disease state markers from one population of subjects, and then validated on an independent group of individuals with MCI, increasing its generalizability. In addition, because the 4-marker signature is in the form of simple decision cut-points, it can readily be applied for clinical trial patient enrollment and in clinical practice for physicians without the need for complex calculations. It will be beneficial in the future to evaluate the performance of this signature in databases containing other neurodegenerative diseases to determine the specificity of these markers against related illnesses. In addition, further evaluation and validation of PTPRN as a diagnostic and progression marker for patients with early signs of cognitive impairment, in conjunction with the core beta-amyloid and tau markers, in line with the ATN construct proposed in the 2018 NIA-AA consensus paper may provide additional insights about AD pathology.

---

**Supplemental Table 1.**
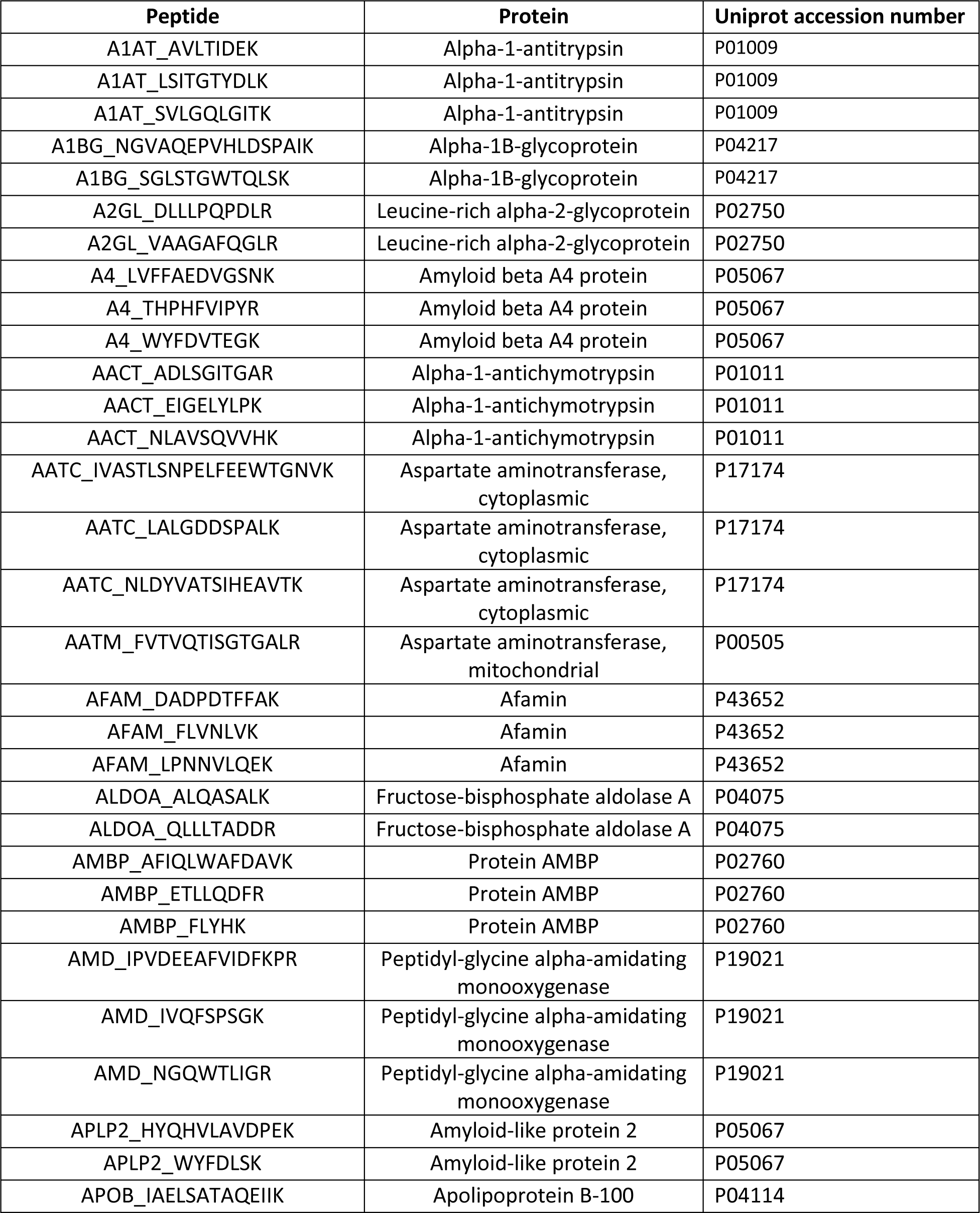
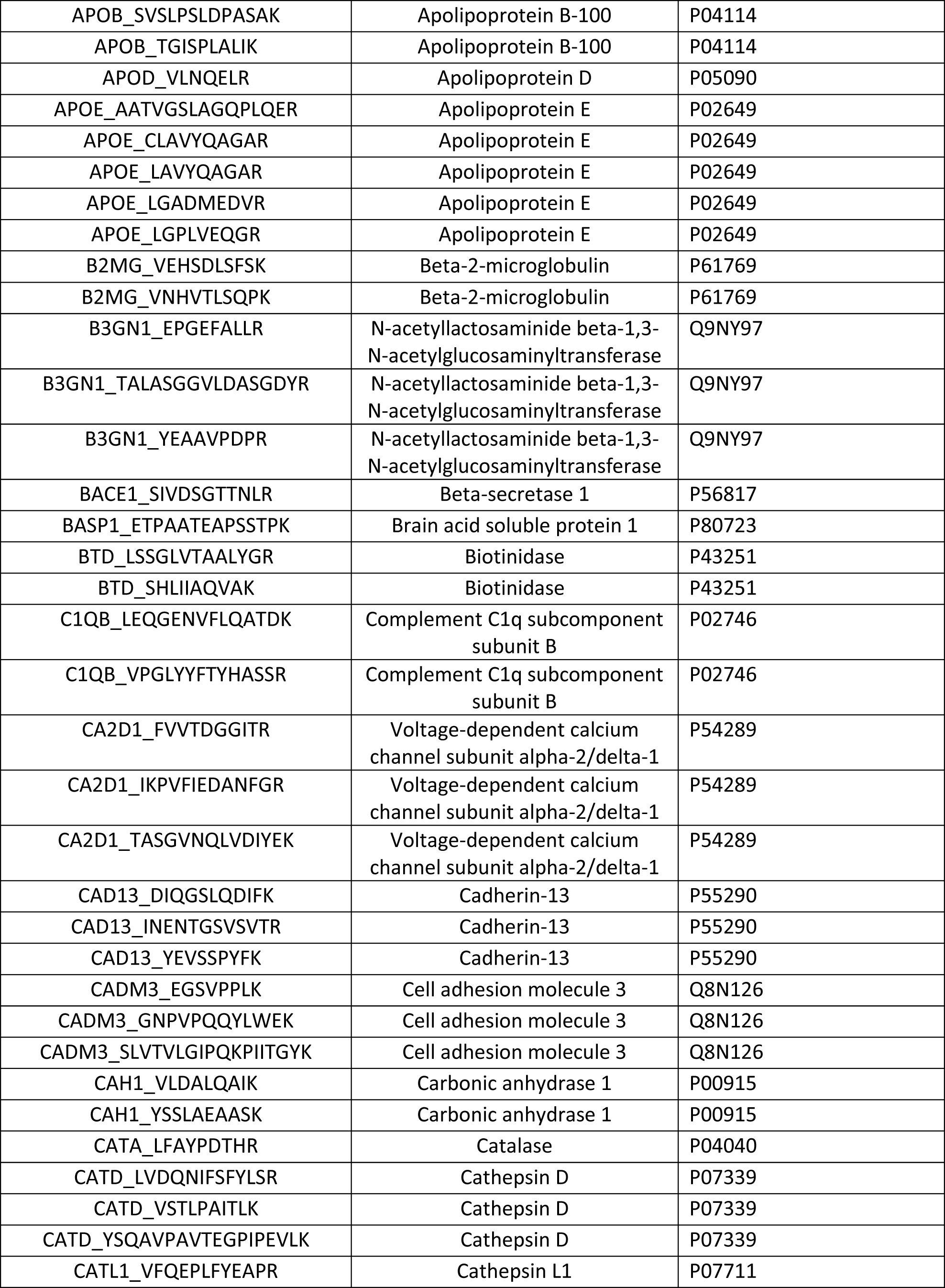
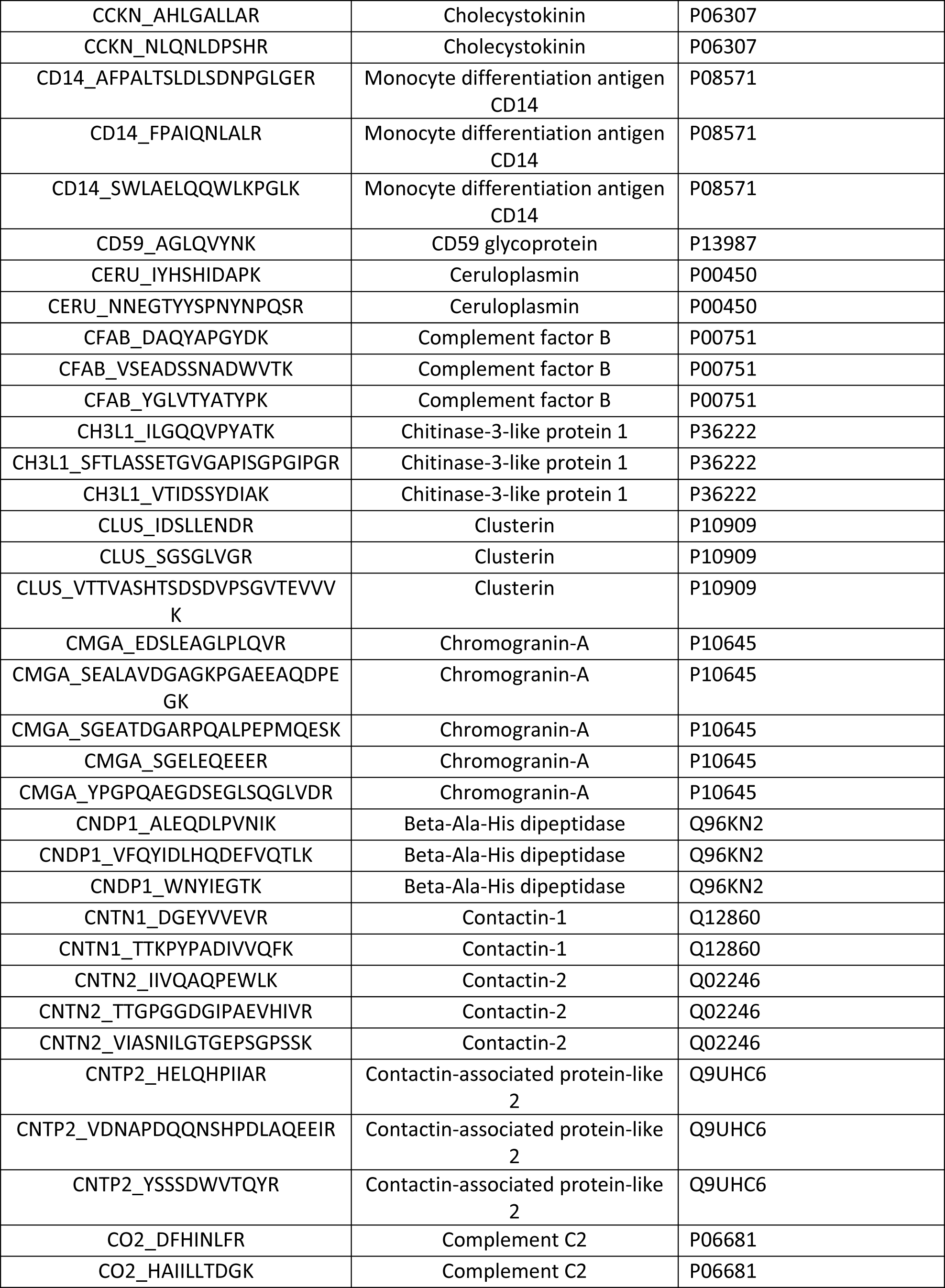
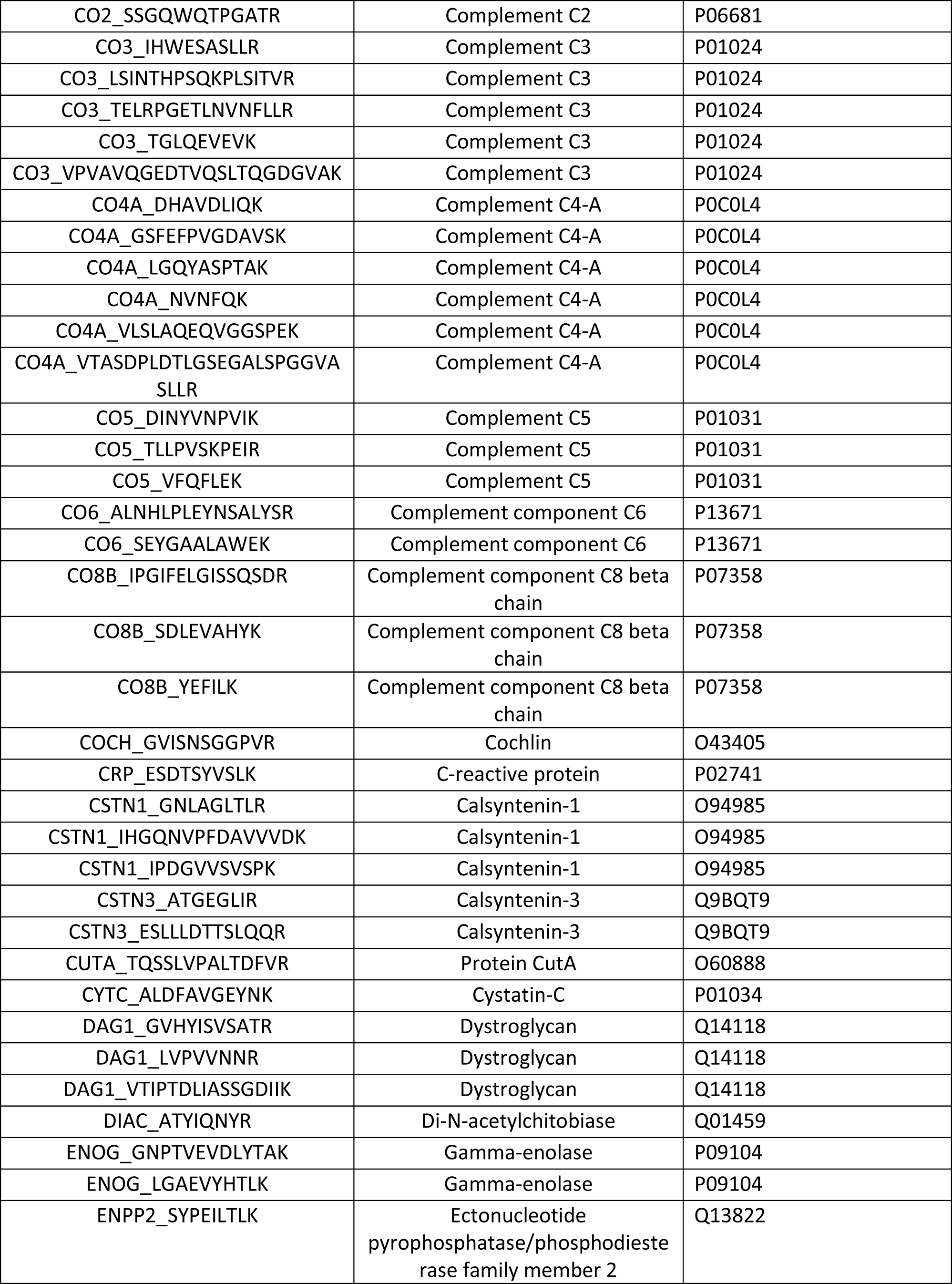
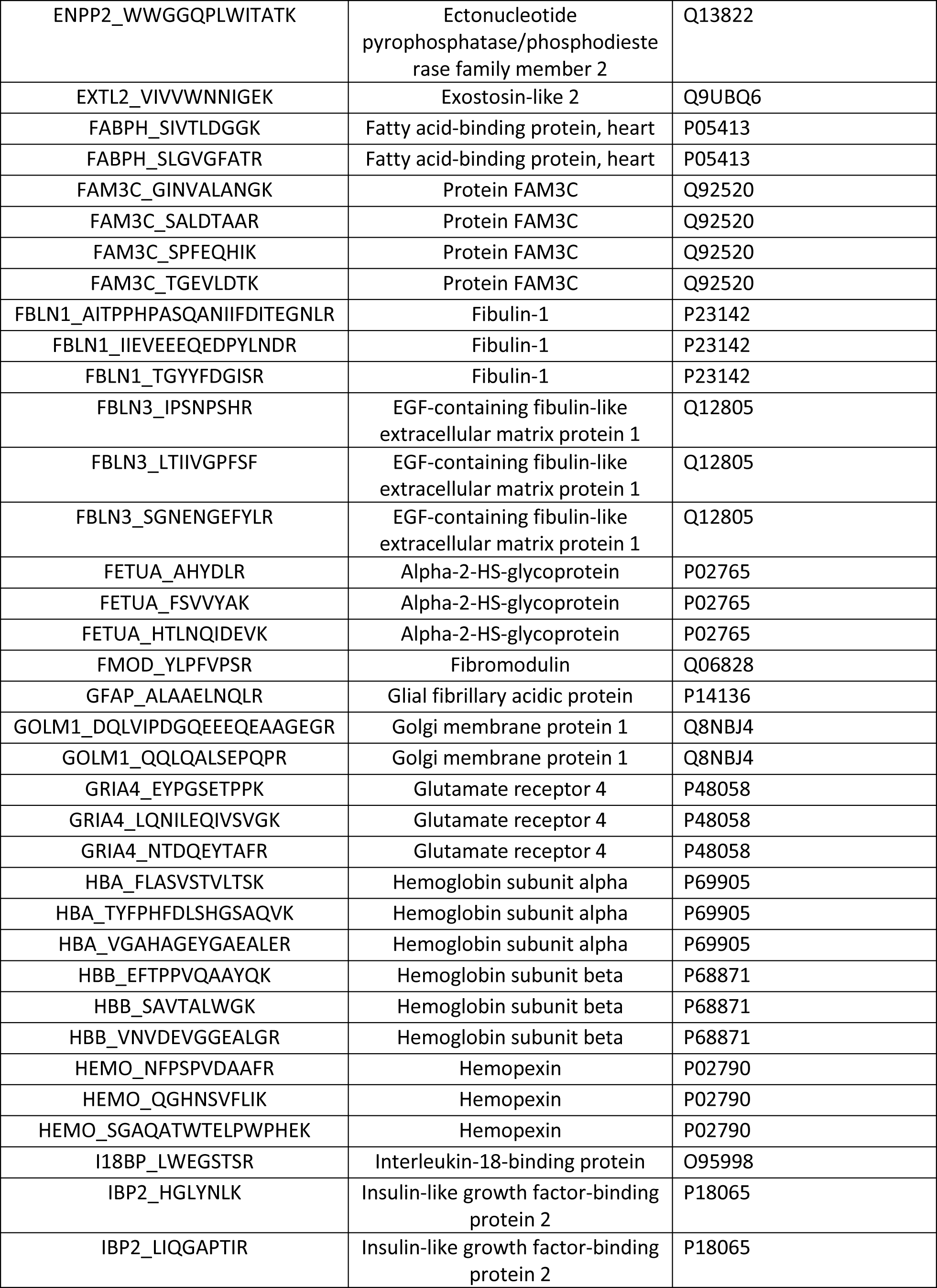
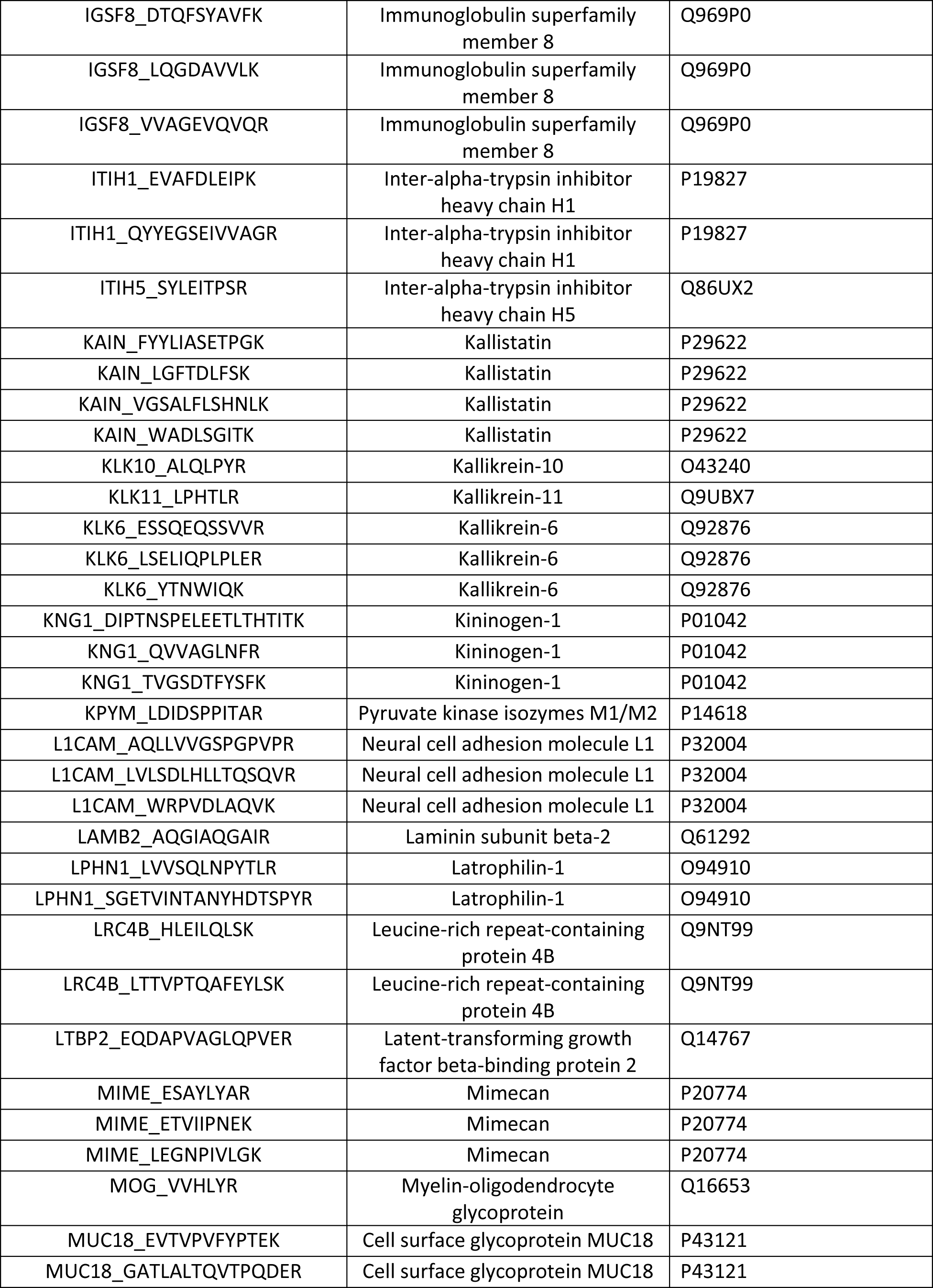
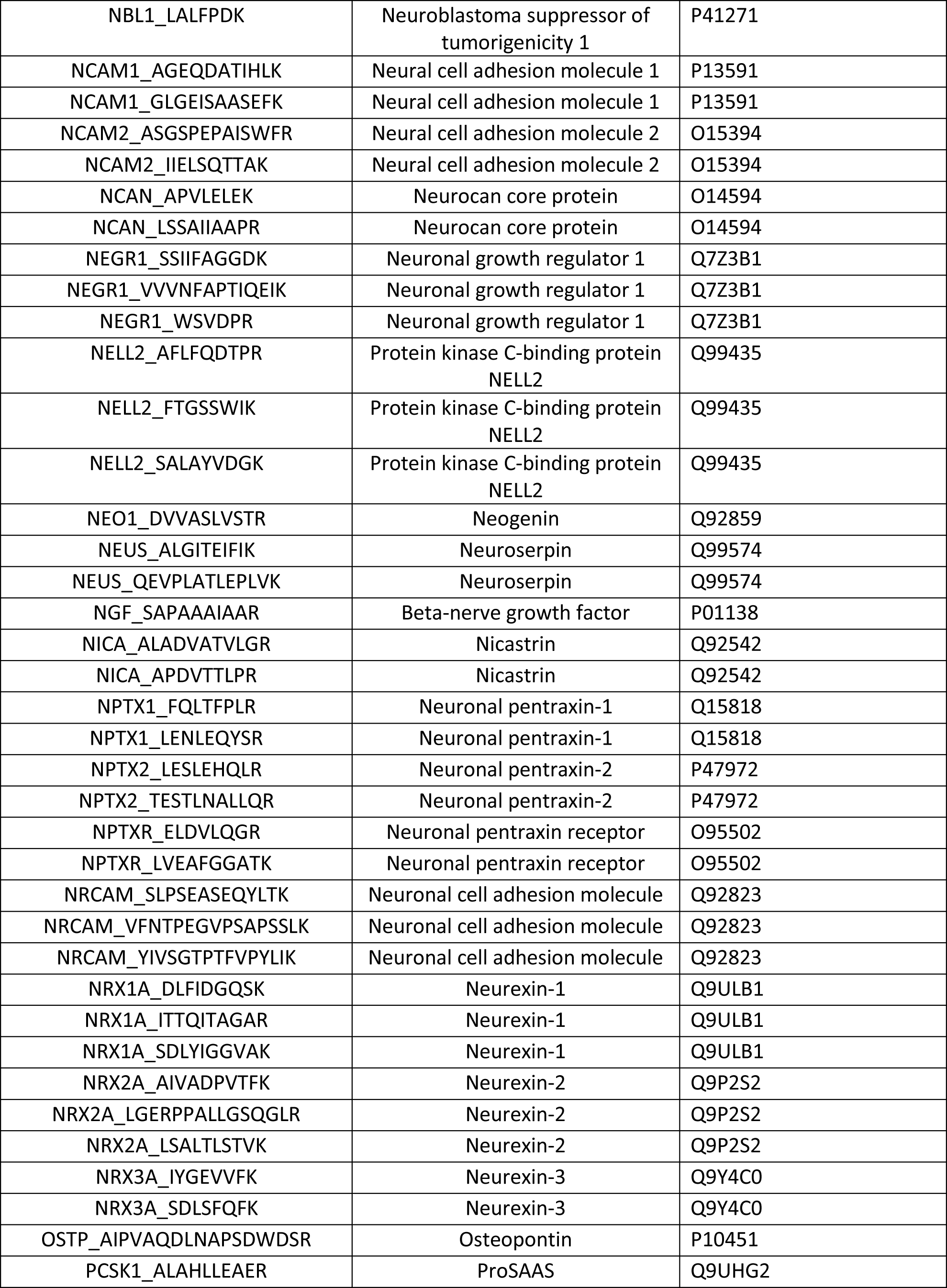
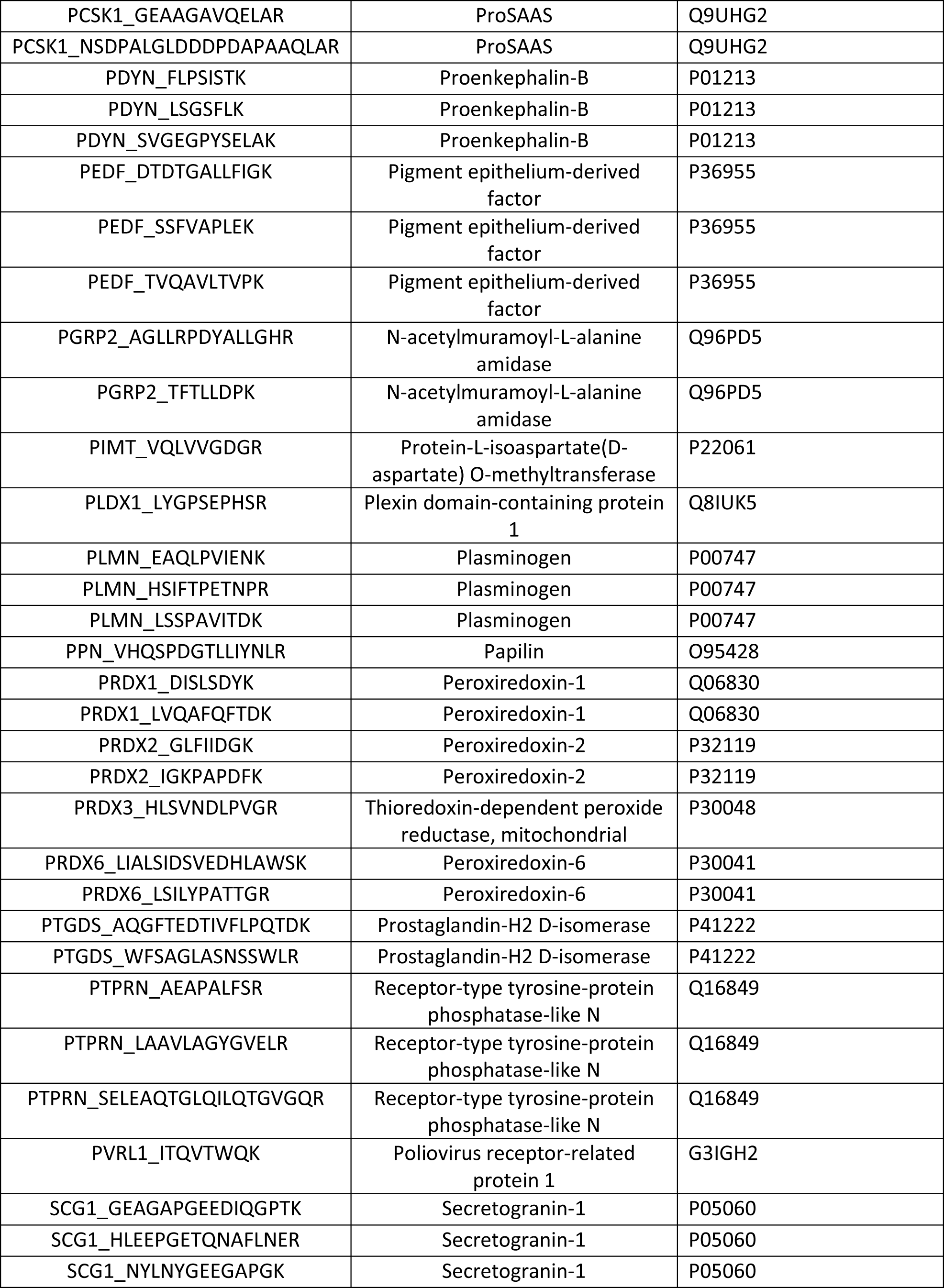
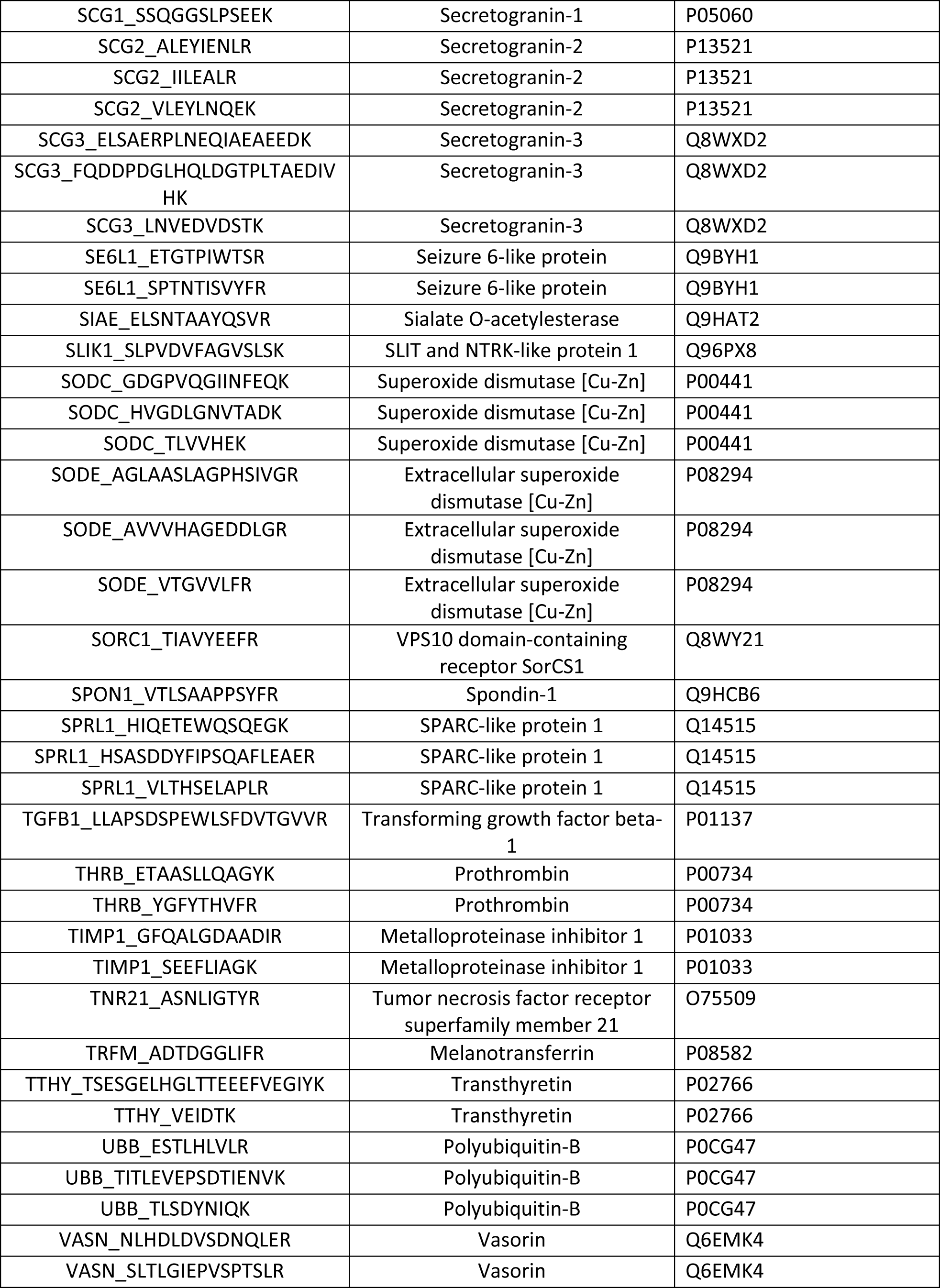
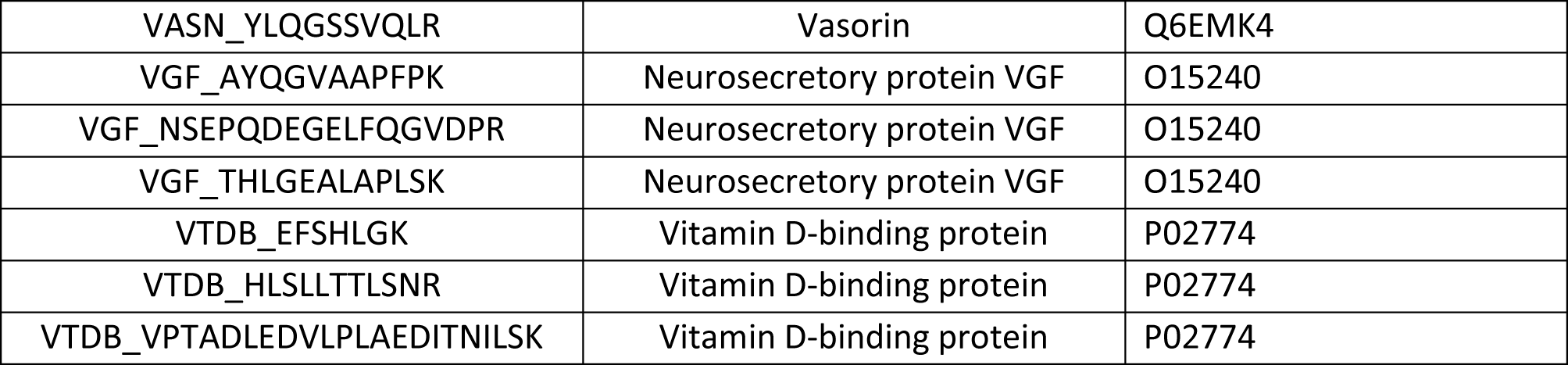
List of Peptides. All peptides, proteins and UniProt accession numbers from the peptides measured in this study.

## Acknowledgements

The authors thank Dr. Clifford Jack for his constructive comments about an earlier version of this manuscript.

## References

[1] Leifer BP (2003) Early diagnosis of Alzheimer’s disease: clinical and economic benefits. Journal of the American Geriatrics Society 51, S281–S288.

[2] Prince M, Bryce R, Ferri C (2011) World Alzheimer Report 2011: The benefits of early diagnosis and intervention, Alzheimer’s Disease International.

[3] Steinberg M, Shao H, Zandi P, Lyketsos CG, Welsh-Bohmer KA, Norton MC, Breitner JC, Steffens DC, Tschanz JT (2008) Point and 5-year period prevalence of neuropsychiatric symptoms in dementia: the Cache County Study. International Journal of Geriatric Psychiatry: A journal of the psychiatry of late life and allied sciences 23, 170–177.

[4] Roque M, Salva A, Vellas B (2013) Malnutrition in community-dwelling adults with dementia (NutriAlz Trial). The journal of nutrition, health & aging 17, 295–299.

[5] Ngandu T, Lehtisalo J, Solomon A, Levälahti E, Ahtiluoto S, Antikainen R, Bäckman L, Hänninen T, Jula A, Laatikainen T (2015) A 2 year multidomain intervention of diet, exercise, cognitive training, and vascular risk monitoring versus control to prevent cognitive decline in at-risk elderly people (FINGER): a randomised controlled trial. The Lancet 385, 2255–2263.

[6] Petersen RC, Smith GE, Waring SC, Ivnik RJ, Tangalos EG, Kokmen E (1999) Mild Cognitive Impairment: Clinical Characterization and Outcome. Arch Neurol 56, 303–308.

[7] Small GW, Kepe V, Ercoli LM, Siddarth P, Bookheimer SY, Miller KJ, Lavretsky H, Burggren AC, Cole GM, Vinters HV (2006) PET of brain amyloid and tau in mild cognitive impairment. New England Journal of Medicine 355, 2652–2663.

[8] Egan MF, Kost J, Tariot PN, Aisen PS, Cummings JL, Vellas B, Sur C, Mukai Y, Voss T, Furtek C (2018) Randomized Trial of Verubecestat for Mild-to-Moderate Alzheimer’s Disease. New England Journal of Medicine 378, 1691–1703.

[9] Ketter N, Brashear HR, Bogert J, Di J, Miaux Y, Gass A, Purcell DD, Barkhof F, Arrighi HM (2017) Central review of amyloid-related imaging abnormalities in two phase III clinical trials of bapineuzumab in mild-to-moderate Alzheimer’s disease patients. Journal of Alzheimer’s Disease 57, 557–573.

[10] Bayer AJ, Bullock R, Jones RW, Wilkinson D, Paterson K, Jenkins L, Millais S, Donoghue S (2005) Evaluation of the safety and immunogenicity of synthetic Aβ42 (AN1792) in patients with AD. Neurology 64, 94–101.

[11] Llano DA, Laforet G, Devanarayan V (2011) Derivation of a New ADAS-cog Composite Using Tree-based Multivariate Analysis: Prediction of Conversion From Mild Cognitive Impairment to Alzheimer Disease. Alzheimer Disease & Associated Disorders 25, 73–84 10.1097/WAD.1090b1013e3181f1095b1098d1098.

[12] Swords GM, Nguyen LT, Mudar RA, Llano DA (2018) Auditory system dysfunction in Alzheimer disease and its prodromal states: A review. Ageing research reviews.

[13] Anderson ED, Wahoske M, Huber M, Norton D, Li Z, Koscik RL, Umucu E, Johnson SC, Jones J, Asthana S (2016) Cognitive variability—A marker for incident MCI and AD: An analysis for the Alzheimer’s Disease Neuroimaging Initiative. Alzheimer’s & Dementia: Diagnosis, Assessment & Disease Monitoring 4, 47–55.

[14] Henriques AD, Benedet AL, Camargos EF, Rosa-Neto P, Nóbrega OT (2018) Fluid and imaging biomarkers for Alzheimer’s disease: Where we stand and where to head to. Experimental gerontology.

[15] Blennow K, Dubois B, Fagan AM, Lewczuk P, de Leon MJ, Hampel H (2015) Clinical utility of cerebrospinal fluid biomarkers in the diagnosis of early Alzheimer’s disease. Alzheimer’s & Dementia 11, 58–69.

[16] Jack CR, Bennett DA, Blennow K, Carrillo MC, Dunn B, Haeberlein SB, Holtzman DM, Jagust W, Jessen F, Karlawish J (2018) NIA-AA Research Framework: Toward a biological definition of Alzheimer’s disease. Alzheimer’s & Dementia 14, 535–562.

[17] Jack CR, Bennett DA, Blennow K, Carrillo MC, Feldman HH, Frisoni GB, Hampel H, Jagust WJ, Johnson KA, Knopman DS (2016) A/T/N: an unbiased descriptive classification scheme for Alzheimer disease biomarkers. Neurology 87, 539–547.

[18] Sunderland T, Linker G, Mirza N, Putnam KT, Friedman DL, Kimmel LH, Bergeson J, Manetti GJ, Zimmermann M, Tang B, Bartko JJ, Cohen RM (2003) Decreased {beta}- Amyloid1-42 and Increased Tau Levels in Cerebrospinal Fluid of Patients With Alzheimer Disease. JAMA 289, 2094–2103.

[19] Shaw LM, Vanderstichele H, Knapik-Czajka M, Clark CM, Aisen PS, Petersen RC, Blennow K, Soares H, Simon A, Lewczuk P (2009) Cerebrospinal fluid biomarker signature in Alzheimer’s disease neuroimaging initiative subjects. Annals of neurology 65, 403–413.

[20] Llano DA, Bundela S, Mudar RA, Devanarayan V, Initiative AsDN (2017) A multivariate predictive modeling approach reveals a novel CSF peptide signature for both Alzheimer’s Disease state classification and for predicting future disease progression. PloS one 12, e0182098.

[21] Chen G, Zhong H, Belousov A, Devanarayan V (2015) A PRIM approach to predictive-signature development for patient stratification. Statistics in medicine 34, 317–342.

[22] Huang X, Sun Y, Trow P, Chatterjee S, Chakravartty A, Tian L, Devanarayan V (2017) Patient subgroup identification for clinical drug development. Statistics in medicine 36, 1414–1428.

[23] Dale AM, Fischl B, Sereno MI (1999) Cortical surface-based analysis: I. Segmentation and surface reconstruction. Neuroimage 9, 179–194.

[24] Fischl B, Sereno MI, Dale AM (1999) Cortical surface-based analysis: II: inflation, flattening, and a surface-based coordinate system. Neuroimage 9, 195–207.

[25] Fischl B, Sereno MI, Tootell RB, Dale AM (1999) High-resolution intersubject averaging and a coordinate system for the cortical surface. Human brain mapping 8, 272–284.

[26] Spellman DS, Wildsmith KR, Honigberg LA, Tuefferd M, Baker D, Raghavan N, Nairn AC, Croteau P, Schirm M, Allard R (2015) Development and evaluation of a multiplexed mass spectrometry based assay for measuring candidate peptide biomarkers in Alzheimer’s Disease Neuroimaging Initiative (ADNI) CSF. Proteomics-Clinical Applications.

[27] Friedman J, Hastie T, Tibshirani R (2010) Regularization paths for generalized linear models via coordinate descent. Journal of statistical software 33, 1.

[28] Efron B, Tibshirani RJ (1994) An introduction to the bootstrap, CRC press.

[29] Shi L, Campbell G, Jones W, Campagne F, Consortium aM (2010) The MicroArray Quality Control (MAQC)-II study of common practices for the development and validation of microarray-based predictive models. Nature biotechnology 28, 827–838.

[30] Corder E, Saunders A, Strittmatter W, Schmechel D, Gaskell P, Small G, Roses A, Haines J, Pericak-Vance MA (1993) Gene dose of apolipoprotein E type 4 allele and the risk of Alzheimer’s disease in late onset families. Science 261, 921–923.

[31] Cai WW, Wang L, Chen Y (2010) Aspartyl aminopeptidase, encoded by an evolutionarily conserved syntenic gene, is colocalized with its cluster in secretory granules of pancreatic islet cells. Bioscience, biotechnology, and biochemistry 74, 2050- 2055.

[32] Uhlén M, Fagerberg L, Hallström BM, Lindskog C, Oksvold P, Mardinoglu A, Sivertsson Å, Kampf C, Sjöstedt E, Asplund A (2015) Tissue-based map of the human proteome. Science 347, 1260419 https://www.proteinatlas.org/ENSG00000054356-PTPRN/tissue/primary+data

[33] Xie J, Zhang B, Lan MS, Notkins AL (1998) Genomic structure and promoter sequence of the insulin-dependent diabetes mellitus autoantigen, IA-2 (PTPRN). Genomics 54, 338–343.

[34] Saeki K, Zhu M, Kubosaki A, Xie J, Lan MS, Notkins AL (2002) Targeted disruption of the protein tyrosine phosphatase-like molecule IA-2 results in alterations in glucose tolerance tests and insulin secretion. Diabetes 51, 1842–1850.

[35] Nishimura T, Kubosaki A, Ito Y, Notkins AL (2009) Disturbances in the secretion of neurotransmitters in IA-2/IA-2β null mice: changes in behavior, learning and lifespan. Neuroscience 159, 427–437.

[36] Kuusisto J, Koivisto K, Mykkänen L, Helkala E-L, Vanhanen M, Hänninen T, Kervinen K, Kesäniemi YA, Riekkinen PJ, Laakso M (1997) Association between features of the insulin resistance syndrome and Alzheimer’s disease independently of apolipoprotein E4 phenotype: cross sectional population based study. Bmj 315, 1045–1049.

[37] Matsuzaki T, Sasaki K, Tanizaki Y, Hata J, Fujimi K, Matsui Y, Sekita A, Suzuki S, Kanba S, Kiyohara Y (2010) Insulin resistance is associated with the pathology of Alzheimer disease The Hisayama Study. Neurology 75, 764–770.

[38] Schrijvers EM, Witteman J, Sijbrands E, Hofman A, Koudstaal PJ, Breteler M (2010) Insulin metabolism and the risk of Alzheimer disease The Rotterdam Study. Neurology 75, 1982–1987.

[39] Steen E, Terry BM, J Rivera E, Cannon JL, Neely TR, Tavares R, Xu XJ, Wands JR, de la Monte SM (2005) Impaired insulin and insulin-like growth factor expression and signaling mechanisms in Alzheimer’s disease–is this type 3 diabetes? Journal of Alzheimer’s disease 7, 63–80.

[40] Kandimalla R, Thirumala V, Reddy PH (2017) Is Alzheimer’s disease a type 3 diabetes? A critical appraisal. Biochimica et Biophysica Acta (BBA)-Molecular Basis of Disease 1863, 1078–1089.

[41] Hokama M, Oka S, Leon J, Ninomiya T, Honda H, Sasaki K, Iwaki T, Ohara T, Sasaki T, LaFerla FM (2013) Altered expression of diabetes-related genes in Alzheimer’s disease brains: the Hisayama study. Cerebral cortex 24, 2476–2488.

[42] Antonell A, Lladó A, Altirriba J, Botta-Orfila T, Balasa M, Fernández M, Ferrer I, Sánchez-Valle R, Molinuevo JL (2013) A preliminary study of the whole-genome expression profile of sporadic and monogenic early-onset Alzheimer’s disease. Neurobiology of aging 34, 1772–1778.

[43] Silva AR, Grinberg LT, Farfel JM, Diniz BS, Lima LA, Silva PJ, Ferretti RE, Rocha RM, Jacob Filho W, Carraro DM (2012) Transcriptional alterations related to neuropathology and clinical manifestation of Alzheimer’s disease. PLoS One 7, e48751.

[44] Sun Y, Nilsson M, Salter H (2013) Genetic interaction analysis of Alzheimer’s disease progression using phospho-tau as a covariate. Alzheimer’s & Dementia: The Journal of the Alzheimer’s Association 9, P555–P556.

[45] Jack Jr CR, Wiste HJ, Vemuri P, Weigand SD, Senjem ML, Zeng G, Bernstein MA, Gunter JL, Pankratz VS, Aisen PS (2010) Brain beta-amyloid measures and magnetic resonance imaging atrophy both predict time-to-progression from mild cognitive impairment to Alzheimer’s disease. Brain 133, 3336–3348.

[46] Bouwman F, Schoonenboom S, van Der Flier W, Van Elk E, Kok A, Barkhof F, Blankenstein M, Scheltens P (2007) CSF biomarkers and medial temporal lobe atrophy predict dementia in mild cognitive impairment. Neurobiology of aging 28, 1070–1074.

[47] Davatzikos C, Bhatt P, Shaw LM, Batmanghelich KN, Trojanowski JQ (2011) Prediction of MCI to AD conversion, via MRI, CSF biomarkers, and pattern classification. Neurobiology of aging 32, 2322. e2319–2322. e2327.

[48] Vemuri P, Wiste H, Weigand S, Shaw L, Trojanowski J, Weiner M, Knopman DS, Petersen RC, Jack C (2009) MRI and CSF biomarkers in normal, MCI, and AD subjects: predicting future clinical change. Neurology 73, 294–301.

[49] Nesteruk M, Nesteruk T, Styczyńska M, Mandecka M, Barczak A, Barcikowska M (2016) Combined use of biochemical and volumetric biomarkers to assess the risk of conversion of mild cognitive impairment to Alzheimer’s disease. Folia neuropathologica 54, 369–374.

[50] Frölich L, Peters O, Lewczuk P, Gruber O, Teipel SJ, Gertz HJ, Jahn H, Jessen F, Kurz A, Luckhaus C (2017) Incremental value of biomarker combinations to predict progression of mild cognitive impairment to Alzheimer’s dementia. Alzheimer’s research & therapy 9, 84.

